# Emergence and the spread of the F200Y benzimidazole resistance mutation in *Haemonchus contortus* and *Haemonchus placei* from buffalo and cattle

**DOI:** 10.1101/425660

**Authors:** Qasim Ali, Imran Rashid, Muhammad Zubair Shabbir, Aziz-Ul-Rahman, Kashif Shahzad, Kamran Ashraf, Neil D. Sargison, Umer Chaudhry

## Abstract

Benzimidazoles have been intensively used in the livestock sector, particularly in small ruminants for over 40 years. This has been led to the widespread emergence of resistance in a number of small ruminant parasite species, in particular *Haemonchus contortus*. In many counties benzimidazole resistance in the small ruminants *H. contortus* has become severely compromising its control; but there is a little information on benzimidazole resistance in *H. contortus* infecting buffalo and cattle. Resistance to benzimidazoles have also been reported in the large ruminant parasite, *Haemonchus placei*, but again there is relatively little information on its prevalence. Hence it is extremely important to understand how resistance-conferring mutations emerge and spread in both parasites in the buffalo and cattle host in order to develop the approaches for the recognition of the problem at an early stage of its development. The present study suggests that the F200Y (T<u>A</u>C) mutation is common in *H. contortus*, being detected in 5/7 populations at frequencies between 7 to 57%. Furthermore, 6/10 *H. placei* populations contained the F200Y (T<u>A</u>C) mutation, albeit at low frequencies of between 0.4 to 5%. The phylogenetic analysis suggests that the F200Y (T<u>A</u>C) mutation in *H. contortus* has emerged on multiple occasions in the region, with at least three independent emergence of resistance alleles across the populations. In contrast, the F200Y (T<u>A</u>C) resistance-conferring mutation in *H. placei* is only seen on a single haplotype. A high level of haplotype frequency of the susceptible alleles in the region, suggests that the unique resistance conferring-mutation has spread from a single emergence; likely by anthropogenic animal movement. Overall, these results provide the first clear genetic evidence for the spread of benzimidazoles resistance-conferring mutations to multiple different locations from a single emergence in *H. placei*; while supporting previous small ruminant-based observations of multiple emergence of resistance mutations in *H. contortus*.

## 1. Introduction

Gastrointestinal (GI) parasitic nematodes are the major cause of disease and productivity loss in grazing livestock costing the North American cattle industry alone more than $2 billion per year (Stromberg and Gasbarre, 2006). *Haemonchus placei* is the most important and highly pathogenic GI nematode species infecting cattle. *Haemonchus contortus* is predominantly a parasite of sheep and goats that also infects large ruminants, with a significant economic impact in tropical and sub-tropical regions (Hoberg et al., 2004; Lichtenfels et al., 1994; Lichtenfels *JR*, 1994). The control of GI nematodes is compromised by the emergence of resistance to anthelmintic drugs (Kaplan and Vidyashankar, 2012), including the benzimidazole group, which is routinely used throughout Asia. The mechanism of benzimidazole resistance has been investigated in small ruminant parasites and strong evidence exists that three different single amino acid substitutions [i.e., F200Y (T<u>A</u>C), F167Y (T<u>A</u>C) and E198A (G<u>C</u>A)] in the isotype-1 β-tubulin are responsible for benzimidazole resistance (Kwa et al., 1994), but there have been few studies of these loci in large ruminants (Chaudhry et al., 2015c). Despite a widespread global focus on the development of benzimidazole resistance in GI nematode parasites of small ruminants, until recently little attention has been given to the possibility of resistance developing in large ruminant parasites (Coles, 2002; Jackson et al., 1987; McKenna, 1996). However, benzimidazole resistance is now emerging in large ruminant parasites and represents a serious challenges to the cattle industry worldwide (Gasbarre et al., 2009a; Sutherland and Leathwick, 2011; Wolstenholme et al., 2004). There have been relatively few investigations into the molecular basis of benzimidazole resistance in large ruminant GI nematode parasites, and in many cases there are few indications that genetic determinants are involved (Edwards and Breckenridge, 1988). Winterrowd et al. (2003), Njue and Prichard (2003) and Demeler et al. (2013) demonstrated that benzimidazole resistance in field isolates of *Cooperia oncophora* and *Ostertagia ostertagi* was associated with the F200Y (T<u>A</u>C) and E198A (G<u>C</u>A) mutations in the isotype-1 ß tubulin locus. The F200Y (T<u>A</u>C) and F167Y (T<u>A</u>C) mutations have been associated with benzimidazole resistance in *H. placei* (Brasil et al., 2012; Chaudhry et al., 2015c).

Understanding the nature of adaptive changes that occur in GI nematodes in response to benzimidazole selection pressure may help to shows how the resistance mutations emerge and spread (Chaudhry, 2015). For example, resistance could emerge as a single mutation and then spread through the parasite population by host migration; in this case a single resistance haplotype would sweep through the population as observed for the small ruminant *H. contortus* benzimidazole resistance allele E198A (G<u>C</u>A) (Chaudhry et al., 2015a). Alternatively, benzimidazole resistance mutations could repeatedly emerge at multiple times and then migrate between parasite populations; in this case multiple resistance haplotypes may sweep through the populations, as recently proposed for small ruminant *H. contortus* benzimidazole resistance alleles F200Y (T<u>A</u>C) and F167Y (T<u>A</u>C) (Brasil et al., 2012; Chaudhry et al., 2015a; Redman et al., 2015; Silvestre and Humbert, 2002). Understanding of the emergence and the spread of benzimidazole resistance mutations in large ruminant *H. contortus* and *H. placei* is poor.

In the present study, we explore the frequency of the F200Y (T<u>A</u>C) benzimidazole resistance mutation in the isotype-1 β-tubulin locus of seven field *H. contortus* and ten *H. placei* populations from buffalo and cattle. We use phylogenetic analysis of *H. placei* and *H. contortus* populations to investigate the emergence and spread of resistance haplotypes. We provide genetic evidence for the emergence of the *H. placei* resistance allele from a single mutation; and for multiple independent emergence of the *H. contortus* resistance allele from a recurrent mutation.

## 2. Materials and Methods

### 2.1. Field parasite samples and genomic DNA extraction

Adult *Haemonchus* worms were obtained from the abomasa of three buffalo and nine cattle, immediately following slaughter at six different abattoirs (Lahore, Faisalabad, Sargodha, Sahiwal, Okara and Gujranwala), where a high prevalence of *Haemonchus* was anticipated (Supplementary Table S1). It was described in our recent study (Ali et al., 2018). Adult worms were fixed in 80% ethanol immediately following removal from the host abomasa. The heads of individual worms were dissected and lysed in single 0.5ml tubes containing 40μl lysis buffer and 10mg/ml Proteinase K (New England BioLabs) previously described by Chaudhry et al. (2016b). 1μl of 1:5 dilutions of each neat single worm lysate was used as a PCR template and identical dilutions of lysate buffer, made in parallel, were used as negative controls. To prepare pooled lysates from each worm population having different number of worms, 1μl aliquots of each individual neat adult worm lysate were pooled. 1μl of a 1:20 dilution of pooled lysate was used as PCR template (Chaudhry et al., 2014).

### 2.2. Haemonchus species-specific pyrosequence genotyping

Genotyping of the single nucleotide polymorphism (SNP) at position 24 (P24) of the rDNA ITS-2 region was used to confirm the species identity of *Haemonchus* spp. in the cattle and buffalo parasite populations recently described by Ali et al. (2018). The rDNA ITS-2 region was amplified from individual *Haemonchus* adult worm lysates using a “universal” forward primer complementary to 5.8S rDNA coding sequence and biotin labelled reverse primer complimentary to the 28S rDNA coding region (Chaudhry et al., 2015b). Final PCR conditions were 1X thermopol reaction buffer, 2mM MgSO_4_, 100μM dNTPs, 0.1μM forward and reverse primers and 1.25U Taq DNA polymerase (New England Biolabs). Thermo-cycling parameters were 95°C for 5 minutes followed by 35 cycles of 95°C for 1 minute, 57°C for 1 minute and 72°C for 1 minute with a single final extension cycle of 72°C for 5 minutes. Following PCR amplification of rDNA ITS-2, the SNP at P24 was determined by pyrosequence genotyping using the PryoMark ID system (Biotage, Sweden). The sequencing primer used was Hsq24 (5’-CATATACTACAATGTGGCTA-3’) and the nucleotide dispensation order was CGAGTCACA. Peak heights were measured using the SNP mode in the PSQ 96 single nucleotide position software. Worms were designated as *H. contortus, H. placei* or putative hybrids based on being homozygous <u>A</u>, homozygous <u>G</u> or heterozygous <u>A/G</u> at position 24, respectively (Ali et al., 2018).

### 2.3. Illumina Mi-seq deep amplicon sequencing of isotype-1 β-tubulin locus

Recently, we have developed and validated the Illumina Mi-seq deep amplicon sequencing technology to study benzimidazole resistance in *Teladorsagia circumcincta* laboratory isolates, based on the isotype-1 β-tubulin locus (MacLeay et al., 2018). The resistance status of as low as 0.1% in a sample was detected accurately. In the present study, Illumina Mi-seq deep amplicon sequencing was used first time for the large-scale survey of benzimidazole resistance from the natural field populations. Therefore species identified *H. contortus* and *H. placei* were further used for Illumina Mi-seq deep amplicon sequencing of the isotype-1 β-tubulin locus encompassing parts of exons 4 and 5 including codons F167Y (T<u>T</u>C-T<u>A</u>C), E198A (G<u>A</u>A-G<u>C</u>A) and F200Y (T<u>T</u>C-T<u>A</u>C) and the intervening intron for both *H. contortus* (328bp) and *H. placei* (325bp). Chaudhry et al. (2016a) used the primer set (HcPYR_For, HcPYR_Rev) to amplify the isotype-1 β-tubulin locus, was modified by adding the adapters to allow the successive annealing of subsequent primers. The N is the number of random nucleotides included between the locus specific primers and the adopter sequences to increase the variety of generated amplicon. Four forward (HcPYR_For, HcPYR_For-1N, HcPYR_For-2N, and HcPYR_For-3N) and four reverse primers (HcPYR_Rev, HcPYR_Rev-1N, HcPYR_Rev-2N, HcPYR_Rev-3N) were mixed in equal proportion (Supplementary Table S1).

The primers were further used for PCR under the following conditions: 5X KAPA HiFi Hot START Fidelity buffer, 10mM dDNTs, 10uM forward and reverse adopter primer, 0.5U KAPA HiFi Hot START Fidelity Polymerase (KAPA Biosystems, USA), 13.25ul ddH_2_O and 1ul of worm lysate. The thermocycling conditions of the PCR were 95°C for 2 minutes, followed by 35 cycles of 98°C for 20 seconds, 62°C for 15 seconds, 72°C for 15 seconds and a final extension 72°C for 5 minutes. PCR products were purified with AMPure XP Magnetic Beads (1X) (Beckman coulter, Inc.) using a special magnetic stand and plate in accordance with the protocols described by Beckman coulter, Inc.

After the purification, a second round of PCR was performed by using eight forward and twelve reverse barcoded primers. The primers were used in manner to ensure that the same forward and reverse primer combinations did not occur in different sample. The second round PCR conditions were 5X KAPA HiFi Hot START Fidelity buffer, 10mM dNTPs, 0.5U KAPA HiFi Hot START Fidelity Polymerase (KAPA Biosystems,USA), 13.25ul ddH2O and 2ul of first round PCR product as DNA template. The barcoded forward (N501 to N508) and reverse (N701 to N712) primers (10uM each) were obtained from Illumina MiSeq manual (http://dnatech.genomecenter.ucdavis.edu). The thermocycling condition of the PCR were 98°C for 45 seconds, followed by 7 cycles of 98°C for 20 seconds, 63°C for 20 seconds, and 72°C for 2 minutes. PCR products were purified with AMPure XP Magnetic Beads (1X) according to the protocols described by Beckman coulter, Inc. The pooled library was measured with KAPA qPCR library quantification kit (KAPA Biosystems, USA). The prepared library was then run on an Illumina MiSeq Sequencer using a 500-cycle pair end reagent kit (MiSeq Reagent Kits v2, MS-103-2003) loaded at 9 pmol with addition 10% Phix Contro v3 (Illumina, FC-11-2003). The MiSeq separated all sequences by sample during post-run processing by recognized indices and to generate FASTAQ files.

### 2.4. MiSeq data handling and bioinformatic filtering

MiSeq data was analysed with our own adapted pipeline. Briefly, an NCBI BLASTN search was used to generate consensus sequences of the isotype-1 β-tubulin locus. Consensus sequences were built from FASTA files using Geneious Pro 5.4 software (Drummond AJ, 2012). Those MiSeq data that did not hit with isotype-1 β-tubulin consensus sequences were discarded as artifactual or contaminating sequences. Samples with arbitrarily less than 1000 reads, were considered as failed preparations. Overall, millions of benzimidazole susceptible and resistance reads of isotype-1 β-tubulin locus were generated from MiSeq data sets of *H. contortus* and *H. placei* populations (Table 1). Polymorphisms appearing more than once in the data set were expected to be real, whereas polymorphisms that only occur once are possible artefacts due to sequencing errors. We used a previously described method to test for this (Chaudhry et al., 2015a; Redman et al., 2015), whereby the distribution of the SNPs was plotted along the isotype-1 β-tubulin locus. This result in the generation of thirty-two *H. contortus* and twenty-five *H. placei* unique isotype-1 β-tubulin haplotypes (Supplementary Table S3) from *H. contortus* and *H. placei* populations.

**Table 1:** Allele frequency (%) of SNPs that resulted in an amino acid change at codons F200Y (TTC/T<u>A</u>C), F167Y (TTC/T<u>A</u>C) and E198A (GAA/G<u>C</u>A) in isotype-1 β-tubulin obtained seven *H. contortus* and ten *H.placei* populations from Punjab province of Pakistan.

| Pakistani field Populations | Host | No of worms in each pool | Total no. of susceptible reads (Illumina Miseq) | Total no. of resistant reads (Illumina Miseq) | Total no. of Illumina MiSeq reads | F167Y |  | E198A |  | F200Y |  | Abattoir Location (Province) |
| --- | --- | --- | --- | --- | --- | --- | --- | --- | --- | --- | --- | --- |
|  |  |  |  |  |  | TTC | T<u>A</u>C | GAA | G<u>C</u>A | TTC | T<u>A</u>C |  |
| *H. contortus* |  |  |  |  |  |  |  |  |  |  |  |  |
| Pop1C | Cattle | 20 | 66511 | 15399 | 81910 | 100 | 0 | 100 | 0 | 81.2 | 18.8 | Lahore, Punjab |
| Pop2C | Cattle | 16 | 20696 | 12953 | 33649 | 100 | 0 | 100 | 0 | 61.5 | 38.5 | Gujranwala, Punjab |
| Pop3B | Buffalo | 20 | 60651 | 33077 | 93728 | 100 | 0 | 100 | 0 | 64.7 | 35.3 | Gujranwala, Punjab |
| Pop8C | Cattle | 2 | 8540 | 0 | 8540 | 100 | 0 | 100 | 0 | 100 | 0 | Sahiwal, Punjab |
| Pop9C | Cattle | 8 | 13544 | 1004 | 14548 | 100 | 0 | 100 | 0 | 93 | 7 | Okara, Punjab |
| Pop10C | Cattle | 1 | 14572 | 0 | 14572 | 100 | 0 | 100 | 0 | 100 | 0 | Sargodha, Punjab |
| Pop12B | Buffalo | 9 | 15821 | 21178 | 36999 | 100 | 0 | 100 | 0 | 42.7 | 57.3 | Okara, Punjab |
| *H. placei* |  |  |  |  |  |  |  |  |  |  |  |  |
| Pop2C | Cattle | 3 | 19980 | 0 | 19980 | 100 | 0 | 100 | 0 | 100 | 0 | Gujranwala, Punjab |
| Pop4C | Cattle | 20 | 42060 | 152 | 42212 | 100 | 0 | 100 | 0 | 99.6 | 0.4 | Lahore, Punjab |
| Pop5B | Buffalo | 20 | 29320 | 0 | 29320 | 100 | 0 | 100 | 0 | 100 | 0 | Faisalabad, Punjab |
| Pop6C | Cattle | 8 | 18198 | 0 | 18198 | 100 | 0 | 100 | 0 | 100 | 0 | Faisalabad, Punjab |
| Pop7C | Cattle | 20 | 4769 | 215 | 4984 | 100 | 0 | 100 | 0 | 95 | 5 | Sahiwal, Punjab |
| Pop8C | Cattle | 18 | 73958 | 320 | 74278 | 100 | 0 | 100 | 0 | 99.5 | 0.5 | Sahiwal, Punjab |
| Pop9C | Cattle | 11 | 76427 | 290 | 76717 | 100 | 0 | 100 | 0 | 99.6 | 0.4 | Okara, Punjab |
| Pop10C | Cattle | 19 | 95522 | 0 | 95522 | 100 | 0 | 100 | 0 | 100 | 0 | Sargodha, Punjab |
| Pop11B | Buffalo | 20 | 104566 | 360 | 104926 | 100 | 0 | 100 | 0 | 99.6 | 0.4 | Sargodha, Punjab |
| Pop12B | Buffalo | 10 | 47288 | 240 | 47528 | 100 | 0 | 100 | 0 | 99.5 | 0.5 | Okara, Punjab |

### 2.5. Split Tree and Medium Joining network analysis of isotype-1 β-tubulin haplotypes

Phylogenetic network analysis of isotype-1 β-tubulin haplotypes has been previously described by Chaudhry et al. (2015a). Briefly, circular Split Tree networks (equal angle) of isotype-1 β-tubulin haplotypes based on genetic distance were generated using the median-net method employed in SplitsTrees4 software (Huson and Bryant, 2006). A Median Joining network of isotype-1 β-tubulin sequences containing all possible shortest trees was generated by setting the epsilon parameter equal to the greatest weighted distance (epsilon = 10) in Network 4.6.1 software (Fluxus Technology Ltd.). The program jModeltest 12.2.0 was used to select the appropriate model of nucleotide substitutions for ML analysis (Posada, 2008). According to the Bayesian information criterion, the best scoring was Hasegawa-Kishino-Yano (HKY+G). The model of substitution was used with parameters estimated from the data. Branch supports were obtained by 1000 bootstraps of the data. The most probable ancestral node was determined by rooting the *H. placei* sequence networks to the closely *H. contortus* outgroup, and *vice versa*.

## 3. Results

### 3.1. H. contortus and H. placei co-infections are common in buffalo and cattle

A total 228 individual *Haemonchu*s worms were pyrosequence genotyped for the rDNA ITS-2 P24 SNP (168 worms form cattle and 60 worms from buffalo) previously described by Ali et al. (2018). In Brief, 76 worms were identified as *H. contortus* (P24 <u>A</u> genotype) and 149 worms were identified as *H. placei* (P24 <u>G</u> genotype) (Supplementary Table S2). All worms were identified as *H. contortus* (homozygous <u>A</u> at rDNA ITS-2 P24) in populations Pop1C of cattle and Pop3B of buffalo and all worms were identified as *H. placei* (homozygous <u>G</u> at rDNA ITS-2 P24) in populations Pop4C, Pop5C, Pop6C and Pop7C of cattle, and Pop11B of buffalo. The remaining populations Pop2C, Pop8C, Pop9C and Pop10C of cattle, and Pop12C of buffalo contained a mixture of *H. contortus* (homozygous <u>A</u> at rDNA ITS-2 P24) and *H. placei* (homozygous <u>G</u> at rDNA ITS-2 P24) indicating co-infection with the two species (Supplementary Table S2). Two cattle and one buffalo host, each co-infected with *H. contortus* and *H. placei*, also contained a single worm with a heterozygous <u>A/G</u> genotype at the rDNA ITS-2 P24 position, suggestive of *H. contortus* / *H. placei* hybrids (Ali et al., 2018).

### 3.2. Isotype-1 β-tubulin locus F167Y, E198A, F200Y SNPs allele frequencies

325bp fragments of *H. placei* and 328bp fragments of *H. contortus* isotype-1 β-tubulin locus was amplified by adapter/barcoded PCR to identify the resistance at positions F167Y (T<u>A</u>C), E198A (G<u>C</u>A) and F200Y (T<u>A</u>C) (Table 1). Benzimidazole resistance-associated SNP F200Y (T<u>A</u>C)was found in five *H. contortus* populations at frequency between 7 and 57% and in six *H. placei* populations with the F200Y (TAC) mutation at frequency between 0.4 and 5% (Table 1). The benzimidazole resistance associated SNPs [E198A (GCA) and F167Y (TAC)] were not detected in any of the populations.

### 3.3. Haplotype distribution of isotype-1 β-tubulin locus

Thirty-two different haplotypes of the *H. contortus* isotype-1 β-tubulin locus were identified among all seven populations. Based on the analysis of each population separately, most contained multiple different haplotypes. Five populations (Pop1C, Pop2C, Pop3B, Pop 9C and Pop12B) comprising of between 8 and 20 worms contained between 1 and 5 F200Y (T<u>A</u>C) benzimidazole resistance-conferring haplotypes, while the benzimidazole resistance-conferring haplotype was absent from the two smallest *H. contortus* populations (Pop8C and Pop10C) comprising of only 1 and 2 worms respectively (Supplementary Table S3). In contrast, there were twenty-five different haplotypes of the *H. placei* isotype-1 β-tubulin locus among ten *H. placei* populations. Based on the analysis of each population separately, most contained single haplotype. The F200Y (T<u>A</u>C) resistance-conferring polymorphism was detected on a single haplotype in six populations (Pop4C, Pop7C, Pop8C, Pop9C, Pop11B and Pop12B) comprising of between 11 and 20 worms (Supplementary Table S3).

### 3.4. Phylogenetic analysis of the isotype-1 β-tubulin locus

In the case of *H. contortus*, Split Tree and the Median Joining networks were generated with all thirty-two distinct haplotypes of the isotype-1 β-tubulin locus (Figs. 1 and 2). The Split Tree analysis showed seven resistant haplotypes (HR1, HR3, HR4, HR6, HR7, HR8, HR9) of the F200Y (T<u>A</u>C) mutation located in three distinct parts of the network (Fig. 1A). There was a higher degree of haplotypic diversity for the susceptible haplotypes compared to the resistance haplotypes, consistent with the F200Y (T<u>A</u>C) SNP being under selection. Nonetheless the F200Y (T<u>A</u>C) resistance SNP was present on haplotypes located in three different parts of the network. Each of these resistance haplotypes were more closely related to one or more susceptible haplotypes indicating that they were multiple independently emergence (Fig. 1). The Median Joining network analysis showed that most of the populations (Pop12B, Pop1C, Pop9C, Pop3B, Pop2C) contained two resistance haplotypes (HR4, HR7) of F200Y (T<u>A</u>C) mutation. The other two resistance associated haplotypes (HR9, HR5) of the F200Y (T<u>A</u>C) mutation were present in three populations (Pop12B, Pop3B, Pop2C) shown in Fig. 2A. The only exceptions were populations (Pop1C, Pop3B) in which a single resistance haplotype (HR1, HR8, HR6, HR3) of F200Y (T<u>A</u>C) was identified (Fig. 2A). The frequency histograms showed that three resistant haplotypes (HR7, HR4, HR1) contained a high frequency of F200Y (T<u>A</u>C) resistance mutations in three populations (Pop2C, Pop3C, Pop12B) and five resistant haplotypes (HR9, HR3, HR8, HR6, HR5) confined a low frequency of F200Y (T<u>A</u>C) resistance mutations in two populations (Pop2C, Pop3C) (Supplementary Fig. 1A).

**Fig. 1.**
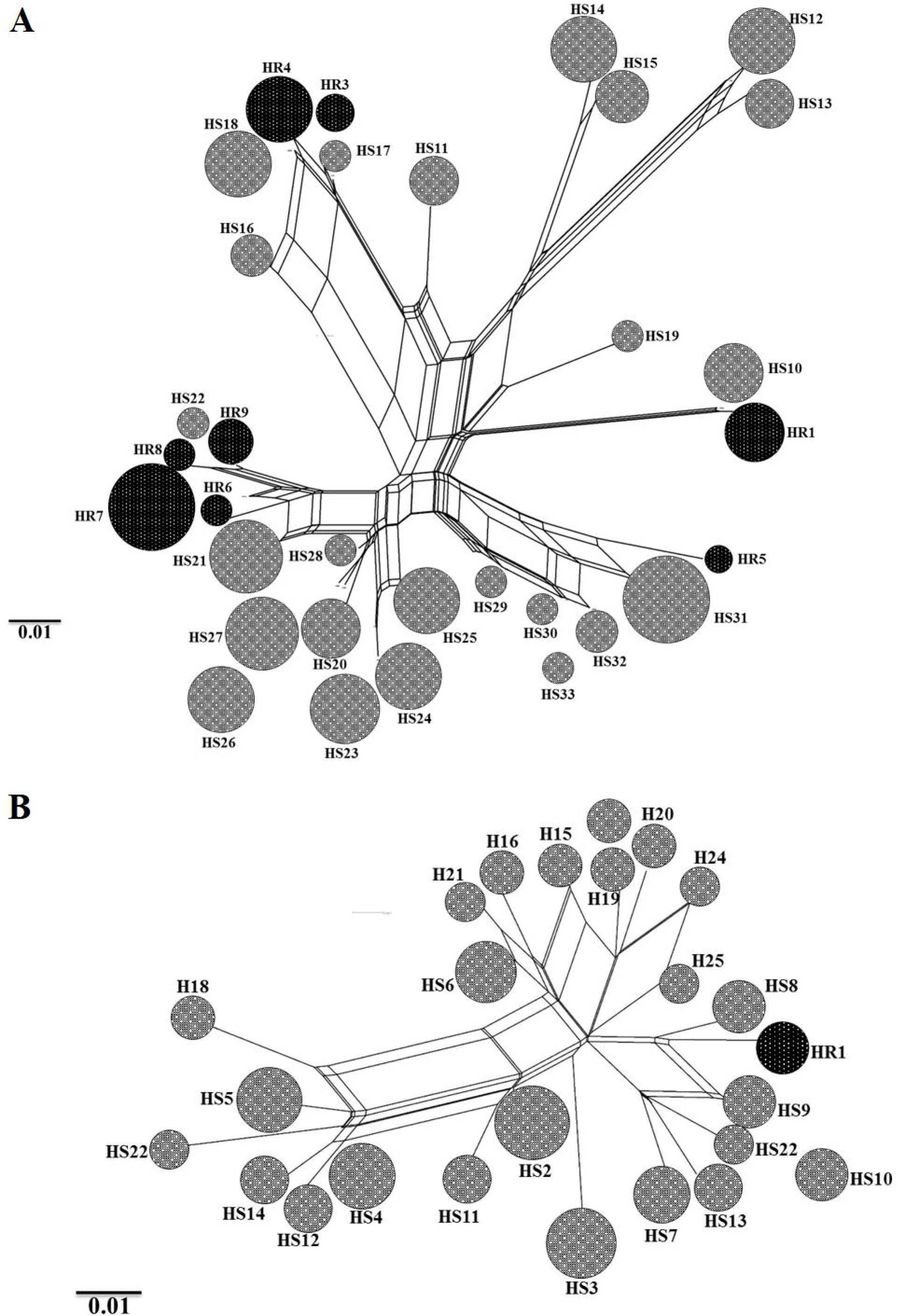
**A & B.** Split Trees network of the *H. contortus* and *H. placei* isotype-1 β-tubulin locus generated with the neighbour-net method of SplitsTrees4 software (Huson and Bryant, 2006). The circles in network represent the different haplotypes and the size of the circles is proportional to the frequency in the population. The haplotypes containing the different mutations are shaded as follows: susceptible haplotypes containing F200Y (TTC)/F167Y (TTC)/E198A (GAA) mutations are light hatch; resistant haplotypes containing F200Y (T<u>A</u>C) mutation is black hatch.

**Fig. 2.**
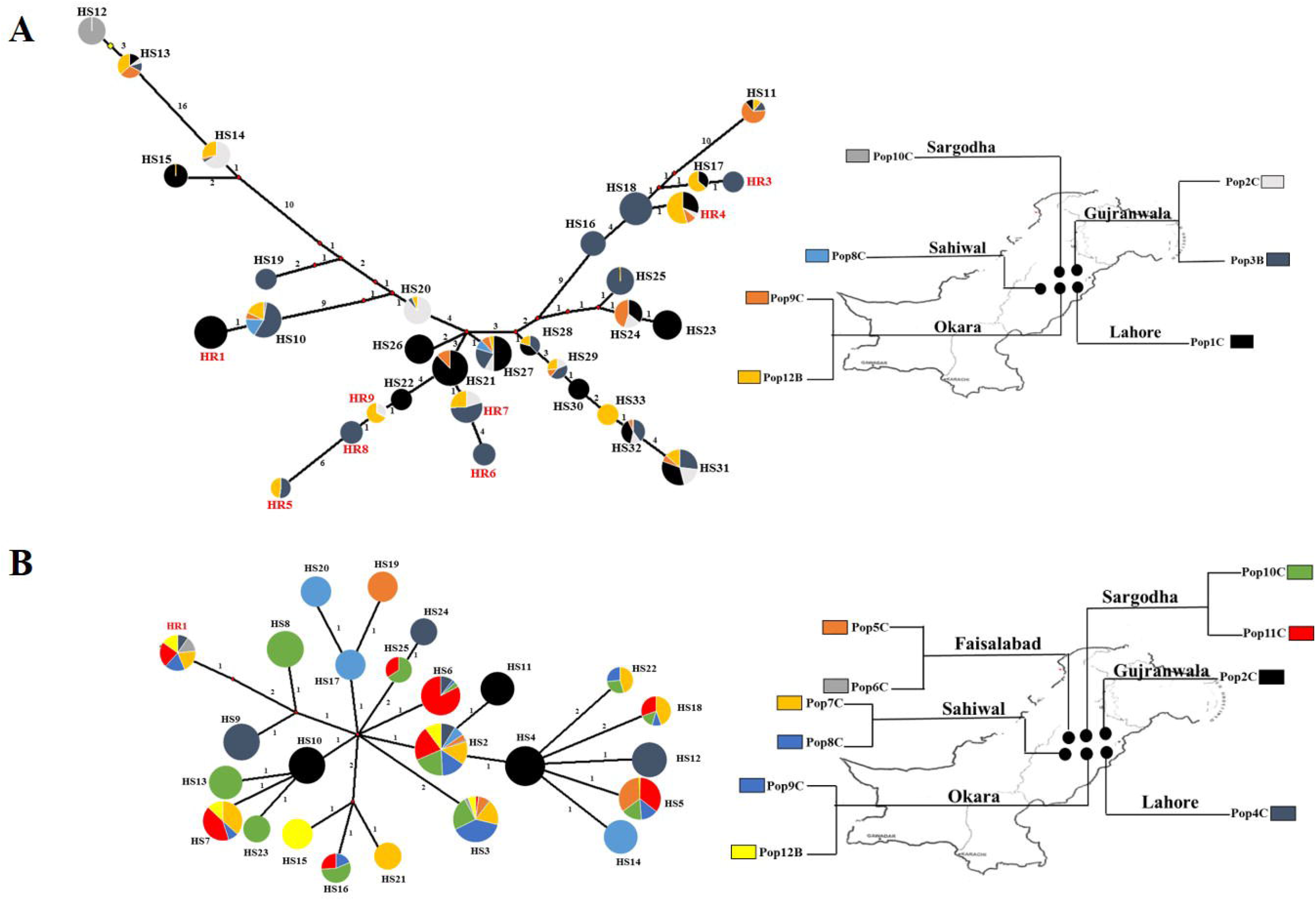
**A & B.** Median Joining network of the *H. contortus* and *H. placei* isotype-1 β-tubulin locus generated in the Network 4.6.1 software (Fluxus Technology Ltd.). A full median network containing all possible shortest trees was generated by setting the epsilon parameter equal to the greatest weighted distance (epsilon = 10). All unnecessary median vectors and links are removed with the MP option (Polzin and Daneschmand, 2003). The size of circle representing each haplotype is proportional to its frequency in the dataset and the colours in the circles reflect the frequency distribution in each population as indicated on the colour key on the inset map. The text providing the name of each haplotype is colour coded as follows; susceptible haplotype F200Y (TTC)/F167Y (TTC)/E198A (GAA) is in black text; P200Y resistant haplotype F200Y (T<u>A</u>C) is in red text.

In the case of *H. placei*, Split Tree and the Median Joining network were generated with all twenty-five distinct haplotypes of the isotype-1 β-tubulin locus (Figs. 1 and 2). The Split Tree network reveals that HR1 haplotype possessing the resistance-conferring F200Y (T<u>A</u>C) SNP, located in single distinct part of the network (Fig. 1B). Although there was also a higher degree of haplotypic diversity seen in the susceptible haplotypes, Nevertheless the F200Y (T<u>A</u>C) resistance SNP was present on single (HR1) haplotype located in single parts of the network. This resistance haplotype was more closely related to one or more susceptible haplotypes indicating that they were single emergence of this mutation (Fig. 1B). The Median Joining network analysis presented that six populations (Pop4C, Pop7C, Pop8C, Pop9C, Pop11B and Pop12B) showed the F200Y (T<u>A</u>C) resistance-conferring SNP on a single haplotype (HR1) (Fig. 2B). The frequency histograms showed that single F200Y (T<u>A</u>C) resistance-associated haplotype (HR1) at law frequency in the dataset was present in six populations (Supplementary Fig. 1B).

## 4. Discussion

The benzimidazoles are one of the most important broad spectrum anthelmintic drug class available for the control of parasitic nematodes of domestic animals and humans (Waller, 1997). They have been intensively used in the livestock sector, particularly in small ruminants, since their first introduction in 1961 (Brown et al., 1961). This has led to the widespread emergence of resistance in a number of small ruminant parasite species including *H. contortus* (Drudge et al., 1964; Smeal et al., 1968). In many developed counties benzimidazole resistance has become so common that the use of this drug class for the control of Haemonchosis is severely compromised (McKellar and Jackson, 2004). There have been numerous studies on the development of the known resistance mutations F200Y, E198A and F167Y in *H. contortus* of small ruminants (Geary et al., 1992; Kwa et al., 1994; Silvestre and Cabaret, 2002), but there has been relatively little research on the same parasite in large ruminants. Additionally, there have been few studies of benzimidazole resistance in the closely related parasite, *H. placei*, hence little is known about the genetics of benzimidazole resistance in this species. However, it was considered likely that resistance would be at an earlier stage of emergence in *H. placei* than in *H. contortus* due to the lower amount of benzimidazole use in cattle than in sheep and the lack of clinical reports of poor treatment efficacy (Yazwinski TA1, 2013). The focus of this work was to investigate the molecular genetics of the known benzimidazole resistance mutations in *H. contortus* and *H. placei* of the large ruminants in the Punjab province of Pakistan. It was anticipated that benzimidazole resistance would be at an earlier stage of development in this region due to less intense selection pressure arising from lower levels of drug use. Consequently, investigating the molecular genetics of benzimidazole resistance in Pakistan not only addresses a major practical knowledge gap, but might also provide new insights into the emergence and spread resistance mutations in parasite populations.

In the present study, first we have demonstrated the frequency of known benzimidazole resistance–conferring mutations F167Y (T<u>A</u>C), E198A (G<u>C</u>A) and F200Y (T<u>A</u>C) in *H. contortus* populations of buffalo and cattle. The F200Y (T<u>A</u>C) mutation was found in two buffalo and three cattle *H. contortus* populations at frequency between 7 and 57%. The resistance has been investigated in the small ruminant parasite *H. contortus*, providing strong evidence exists that three different SNPs [i.e., F200Y (T<u>A</u>C), F167Y (T<u>A</u>C) and E198A (G<u>C</u>A)] in the isotype-1 β-tubulin are responsible for benzimidazole resistance (Brasil et al., 2012; Ghisi et al., 2007; Hoglund et al., 2009; Kotze, 2012; Kwa et al., 1994; Redman et al., 2015; Rufener et al., 2009; Silvestre and Cabaret, 2002; Silvestre and Humbert, 2002). The F200Y (T<u>A</u>C) is the predominant mutation causing widespread, albeit low frequency benzimidazole resistance in sub-tropical developing countries including India and Pakistan (Chaudhry et al., 2015a; Chaudhry et al., 2016b). For example, the F200Y (TAC) mutation was only detected at a very low frequency of between 2 to 6% in *H. contortus* from small ruminants sourced from different rural locations of Pakistan (Chaudhry et al., 2016b). In contrast, high frequencies of the F200Y (T<u>A</u>C) resistance-conferring mutation are widespread in *H. contortus* in developed countries including Australia, New Zealand, France, Sweden, Brazil and UK. This suggests that this polymorphism has little fitness cost, at least in the presence of drug selection (Bisset SA et al., 2014; Brasil et al., 2012; Hoglund et al., 2009; Kotze et al., 2012; Redman et al., 2015; Silvestre and Humbert, 2002). Although the F167Y (T<u>A</u>C) has been detected in countries including USA, Canada, UK, France and Brazil, it is generally present at lower frequencies than the F200Y (T<u>A</u>C) mutation. This may suggests that this polymorphism has a higher fitness cost than the F200Y (T<u>A</u>C) mutation (Barrère et al., 2012; Barrere et al., 2013a; Barrere et al., 2013b; Brasil et al., 2012; Silvestre and Cabaret, 2002). One notable exception to this appears in a study of UK farms where the F167Y (T<u>A</u>C) mutation was present at a higher frequency then the F200Y (T<u>A</u>C) mutation in *H. contortus* populations of small ruminants (Redman et al., 2015). A similar argument applies to the E198A (G<u>C</u>A) SNP which is even rarer than the F167Y (T<u>A</u>C) mutation. It has been detected in just three field-derived populations of *H. contortus* from South Africa (Rufener et al., 2009), Australia (Ghisi et al., 2007) and India (Chaudhry et al., 2015a). In contrast to knowledge of benzimidazole resistance-conferring SNPs in *H. contortus* from small ruminants, there are no genetic information of benzimidazole resistance in *H. contortus* from large ruminants.

In the present study, we have also confirmed the frequency of known benzimidazole resistance–conferring mutations F167Y (T<u>A</u>C), E198A (G<u>C</u>A) and F200Y (T<u>A</u>C) in *H. placei* populations of buffalo and cattle. We detected the F200Y (T<u>A</u>C) mutation in two buffalo and four cattle *H. placei* populations at frequencies between 0.4% to 5%. These results suggested that F200Y (T<u>A</u>C) resistance mutation is likely to be present in many *H. placei* populations of buffalo and cattle. Previously benzimidazole resistance has been reported in *H. placei* in Brazil (Bricarello et al., 2007), Argentina (Anziani et al., 2004), USA (Gasbarre et al., 2009b; Gasbarre et al., 2009a) and north India (Yadav and Verma, 1997), but there is little information on its prevalence. These reports are based on faecal egg count reduction tests (FECRTs) to give an estimate of the benzimidazole resistance or susceptibility status of a parasite community as a whole, but cannot be used to assess the proportion of benzimidazole resistance worms in a particular population (Achi et al., 2003; Jacquiet et al., 1998). The low frequency presence of the F200Y (T<u>A</u>C) mutation has seen in those studies, would not be expected to result in loss of efficacy of benzimidazole drugs detectable using the FECRT. Although there have been relatively few investigations into the molecular genetics of benzimidazole resistance in cattle GI nematode parasites, there are some indications that same genetic determinants are involved in different species. Winterrowd et al. (2003) amplified partial isotype-1 β-tubulin sequences from *Cooperia oncophora* and demonstrated that benzimidazole resistance in field isolates was associated with the F200Y mutation. Njue and Prichard (2003) and Demeler et al. (2013) subsequently analysed isotype-1 β-tubulin from *C. oncophora* and found a small proportion of individuals that carried the F200Y resistant isotype-1 β-tubulin allele. The F200Y (T<u>A</u>C) and F167Y (T<u>A</u>C) polymorphisms have been previously been identified in *H. placei* (Brasil et al., 2012; Chaudhry et al., 2015c), however there is no previous reports of the presence of E198A (G<u>C</u>A) resistance mutations in *H. placei* under natural field conditions.

The presence of the F200Y (T<u>A</u>C) resistance mutation on eight diverse haplotypes suggests that resistance emerged at least three times in the *H. contortus* populations of two buffalo and three cattle hosts. Within the Split Tree analysis, each of three groups of the eight F200Y (T<u>A</u>C) resistance haplotypes (HR4/HR18/HR3; HR9/HR9/HR6/HR7; HR1/HR5) was more closely related to the large number of diverse susceptible haplotypes (twenty-four different susceptible haplotypes in the dataset). The phylogenetic analysis of *H. contortus* haplotypes, therefore, suggests that the F200Y (T<u>A</u>C) resistance mutation arose at multiple independent times. In this case, it appears that the F200Y (T<u>A</u>C) mutation spread to multiple different locations of the Punjab province of Pakistan through the movement of cattle and buffalo. The Median Joining network analysis supported the view that the P200Y (T**A**C) mutation spread much more commonly in *H. contortus* populations. This is further supported by five populations (Pop12B, Pop1C, Pop9C, Pop3B, Pop2C) having two resistance haplotypes (HR4, HR7) of the F200Y (T<u>A</u>C) mutation, while the other two resistance associated haplotypes (HR9, HR5) of the F200Y (T<u>A</u>C) mutation were present in three populations (Pop12B, Pop3B, Pop2C). Hence our study suggests that the F200Y (T<u>A</u>C) resistance mutation spread across several locations in the Punjab province of Pakistan from multiple emergence. This is consistent with previous findings of multiple independent time emergence of the F200Y (T<u>A</u>C) resistance mutation and subsequent spread by widespread animal movements in small ruminant *H. contortus* (Brasil et al., 2012; Chaudhry et al., 2015a; Chaudhry et al., 2016a).

The F200Y (T<u>A</u>C) resistance mutation is present on a single haplotype suggesting that resistance emerged once in the *H. placei* populations of two buffalo and four cattle host. In contrast, there is a large amount of susceptible haplotype diversity within the fragment sequences (twenty-four different susceptible haplotypes in the dataset). The phylogenetic analysis of *H. placei* haplotypes, therefore, suggests that the F200Y (T<u>A</u>C) resistance mutation emerged rarely and that this mutation then spread to multiple different locations of the Punjab province from a single emergence. Given the high level of susceptible haplotype diversity, it would be extremely unlikely that the F200Y (T<u>A</u>C) repeatedly emerged only on the same haplotype (HR1). This represents the first clear genetic evidence of the single emergence and spread of a benzimidazole resistance mutation across multiple locations in the Punjab province of Pakistan.

There is need for the better understanding of the interactions between parasite epidemiology, farming practices, the emergence and spread of anthelmintic resistance mutations to inform sustainable GI nematode control in different parts of the world. In most countries where benzimidazole resistance is well advanced, the phylogenetic diversity of the resistance mutations and the complexity of their relationships hinders the use of genetic analysis to definitively demonstrate how a particular resistance allele has spread from one location to another (Chaudhry, 2015). However, our study in the Punjab region, where resistance is at an earlier stages, provides a simpler situation from which to draw a conclusion. The multiple emergence of the F200Y (T<u>A</u>C) resistance-conferring mutation in *H. contortus* and its subsequent is clearly shown. The detection of a single F200Y (T<u>A</u>C) haplotype in *H. placei* at multiple sites provides persuasive evidence of a single emergence in the region. The way in which this mutation has become widespread from a single emergence provides a clear illustration of the role of migration in the spread of resistance alleles. This emphasises the critical importance of biosecurity measures and quarantine anthelmintic treatments in managing the emergence of resistance in any country where there is significant animal movement. It also implies that the migration of resistance mutations between locations plays an important role in producing the complex patterns of resistance haplotypes seen at the later stages of resistance development.

## Acknowledgment

We are grateful to the Vice Chancellor (Prof. Dr. Talat Naseer Pasha) of the University of Veterinary and Animal Science Lahore Pakistan for his great support in the organization of collection of samples from abattoirs. The study was financially supported by the Higher Education Commission of Pakistan. We would also like to thank the Moredun Research Institute Scotland for their kind support to use pyro-sequencer. The article is the sequel of Dr Umer Chaudhry thesis, which is available in full at the University of Calgary, Canada website.

## Supplementary Figure Legends

**Supplementary Fig. S1 A & B.** Frequency histograms showing resistant and susceptible isotype-1 β-tubulin haplotypes identified from seven *H. contortus* and ten *H. placei* populations. F200Y (TTC)/F167Y (TTC)/E198A (GAA) susceptible haplotypes are shown in light hatch, F200Y (T<u>A</u>C) resistant haplotypes black hatch.

